# A rapid and tunable method to temporally control Cas9 expression enables the identification of essential genes and the interrogation of functional gene interactions in vitro and in vivo

**DOI:** 10.1101/023366

**Authors:** Serif Senturk, Nitin H. Shirole, Dawid D. Nowak, Vincenzo Corbo, Alexander Vaughan, David A. Tuveson, Lloyd C. Trotman, Adam Kepecs, Frank Stegmeier, Raffaella Sordella

## Abstract

The Cas9/CRISPR system is a powerful tool for studying gene function. Here we describe a method that allows temporal control of Cas9/CRISPER activity based on conditional Cas9 destabilization. We demonstrate that fusing an FKBP12-derived destabilizing domain to Cas9 (DD-CAS9) enables conditional Cas9 expression in vitro in the presence of an FKBP12 synthetic ligand and temporal control of gene-editing. Further, we show that this strategy can be easily adapted to co-express, from the same promoter, DD-Cas9 with any other gene of interest, without the latter being co-modulated. In particular, when co-expressed with inducible Cre-ER^T2^, our system enables parallel, independent manipulation of alleles targeted by Cas9 and traditional recombinase with single-cell specificity. We anticipate this platform will be used for the systematic identification of essential genes and the interrogation of genes functional interactions.

## INTRODUCTION

Temporal, spatial, and locus-specific control of gene expression is essential for understanding gene function in biological systems. While ectopic expression of cDNAs has been possible for several decades, the ablation of gene expression has been more problematic. The use of conditional alleles under control of engineered recombinases, as well as antisense and RNAi-mediated gene silencing, have enabled gene knock-down *in vitro* as well as *in vivo* with varying degrees of success (1-3). More recently new genome-editing methods, such as zinc-finger nucleases (ZFNs), transcription activator-like nucleases (TALENs) and Clustered, regularly interspaced, short palindromic repeats and the Cas9 endonuclease (CRISPR/Cas9), have been shown to be powerful tools for directly mutating the genome through targeted gene deletions (4-6). Among these CRISPR/Cas9 has proven to be both efficient and robust across a variety of genetic contexts (7, 8). In this system, the Cas9 nuclease is directed to specific DNA sequences by an engineered small guide RNA (sgRNA) (6). The resulting cleavage sites are repaired by non-homologous end-joining (NHEJ), with the consequent introduction of random mutations into the targeted gene and functional inactivation of genes at high frequency (6).

One of the main problems limiting the use of current CRISPR systems is the constitutive endonuclease activity when Cas9 and its sgRNA are co-expressed. This can be particularly problematic when targeting genes that are developmentally important or essential for viability. Furthermore, it has been shown that constitutive expression of Cas9 can increase the number of off-target mutations and can trigger a DNA damage response (9, 10).

Hence, we sought to improve upon existing CRISPR/Cas9 techniques to generate a system that (1) would provide potent, robust and temporally controlled gene editing, (2) be applicable to a broad spectrum of cell types and tissues, (3) facilitate high throughput manipulation and (4) be tractable. To this end, we exploited recently developed strategies in which a cell-permeable ligand is used in conjugation with a single genetically encoded destabilizing domain (DD) to regulate the expression of any protein of interest (14). The addition of this DD-tag to any protein partner results in proteasomal degradation of the entire fusion product. Notably, addition of a high affinity ligand stabilizes the DD domains in a dose-dependent manner and thus the fusion protein as a whole (15). This approach enables ligand-mediated control of protein stability and function in a highly specific, reversible and dose-dependent manner. The first DD technology was developed based on the identification of specific destabilizing mutations in the human FKBP12 protein, and the development of a family of highly specific synthetic FKBP12 ligands that are highly cell permeable and nontoxic in cultured cells and animals (14). The small-molecule ligand Shield-1 was shown to bind to mutant FKBP12 more tightly than to wild type by three orders of magnitude (16) and to prevent proteasome-induced degradation of DD-YFP and other fusion proteins (17). This destabilized-domain system has been demonstrated to work well for a variety of proteins including kinases, cell cycle regulatory proteins and small GTPases expressed in cultured cells as well as in mice (14, 18).

By fusing the FKBP12-derived destabilizing domain to Cas9, generating DD-Cas9, we demonstrated that this method of conditional regulation of protein stability could be exploited for rapid and reversible Cas9 expression in vitro.

We validated the efficiency of this new platform by conditionally targeting a variety of genes controlling diverse biological processes such as oncogenic transformation (EGFR), DNA damage responses (Tp53), mitochondria metabolism (CypD) and DNA replication/repair (RPA3). By targeting the RPA3 gene and EGFR in the “EGFR-addicted” cells, in particular, we demonstrated the ability of the system to identify genes that are essential for sustained cell growth and survival.

One important aspect of this method that makes it unique is the conditional regulation of CAS9 protein expression independently of its mRNA expression. Hence, this vector system can be easily adapted to co-express, from the same promoter, DD-Cas9 with any other gene of interest, without the latter being co-modulated. We demonstrated this approach could be exploited for generating tractable systems and for interrogating genes functional interactions in vivo.

Moreover when DD-Cas9 is coupled with a conditional Cre allele (Cre-ER^T2^), this platform could be utilized to facilitate the analysis of genes that modulate disease onset and progression in a variety of pre-existing mouse models of human disease based on Cre-lox system.

In summary, our data indicate that fusing Cas9 to a destabilizing domain provides a highly efficient and potent, easy scalable, robust and tunable new modality for temporal control of gene editing that can be applicable to a broad spectrum of in vitro and in vivo models.

## RESULTS

### Establishment of an Inducible Lentiviral Guide RNA Expression System based on conditionally destabilized Cas9

CRISPR/Cas9 technology has been widely utilized to create heritable changes in the genome. However, mutations often result in cell lethality, functional deficits and developmental defects limiting the utility of such models for studying gene function. Additionally, constitutive expression of Cas9 may result in toxicity and in the generation of off-target effects (19).

To overcome these limitations, we generated a temporally controlled Cas9 expression system by fusing Cas9 to an engineered mutant of the human FKBP12 protein (DD-Cas9). As previously shown, the presence of this destabilizing peptide induces the rapid degradation of the fused-protein by the proteasome system (14). Yet in the presence of a synthetic ligand (shield-1), the peptide as well as the fusion partner are effectively shielded from degradation (Fig. 1a). We developed a dual lentiviral vector system consisting of a sgRNA cassette driven by the constitutive U6 promoter and a second module in which Cas9 is fused at its N-terminal with a ligand-responsive destabilizing domain derived from an engineered FKBP12 mutant (DD-Cas9) under the control of a EFS promoter.

**Figure 1:**
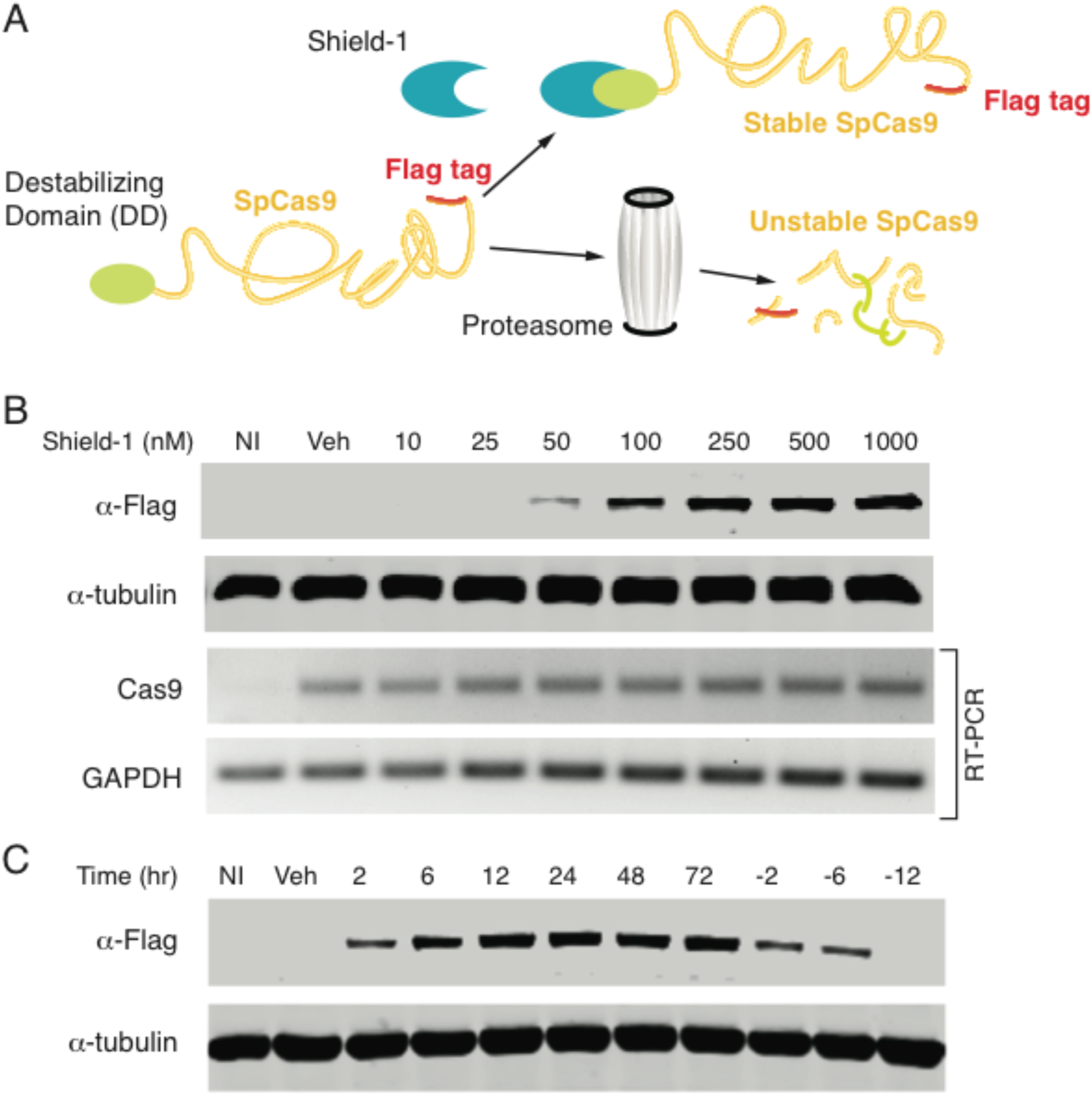
Establishment of an Inducible Lentiviral Guide RNA Expression System based on conditionally destabilized Cas9 (DD-Cas9). **A)** Diagram of conditionally destabilized Cas9. Cas9 is fused to the destabilizing domain of mutant FKB12 that triggers the rapid degradation of the fused-protein. The synthetic ligand (Shield-1) prevents this destabilizing effect and prevents Cas9 from degradation. Of note a Flag-tag its present at the C-terminal of CAS9.**(B)** Ligand-dependent stabilization of DD-Cas9. Cells expressing destabilized Cas9 (DD-Cas9) and uninfected cells (NI) were either mock-treated (vehicle) or treated with increasing concentrations of Shield-1 as indicated. To measure level of expression of Cas9, lysates were analyzed by Western blot with an anti-Flag antibody. Various degrees of Shield-1-induced stabilization could be observed in a dose dependent manner. DD-Cas9 was not detected in lysate from mock-treated cells. To verify that the destabilization of DD-Cas9 occurs post-transcriptionally, DD-Cas9 expression was also verified by RT-PCR analysis using Cas9 specific primers and GAPDH as loading control. **(C)** Rapid and reversible regulation of destabilized DD-Cas9. A549 cells transduced with the DD-Cas9 lentiviral vector and uninfected cells were treated with Shield1 (200 nM) for different time points as indicated. Samples were assayed by western blot analysis at the indicated time points.

To test for the feasibility of this approach, we transduced our engineered lentiviral construct into A549 cells and we measured the levels of CAS9 in the presence and absence of Shield-1 by western blot and RT-PCR analyses (Fig. 1b). Antibodies against Flag were used to indirectly gauge CAS9 levels. Although CAS9 was not detected in uninfected cells or in lysates of mock-treated cells, cells treated with Shield-1 showed strong expression of the expected fusion protein in a dose dependent manner. Importantly, all experimental conditions presented with similar levels of expression of Cas9 mRNA in the presence or absence of synthetic ligand (Fig. 1b).

We next assayed the destabilizing domain for kinetics of protein degradation. Cells were treated with Shield-1 and harvested 2, 6, 12, 24, 48 and 72 h after ligand treatment. Upon just 2 hours of treatment with Shield-1, we observed a rapid induction of CAS9 protein expression compared to uninfected and mock-control cells (Fig. 1c). This effect was reversible upon 2 hours of ligand withdrawal from the media, with protein levels becoming negligible within 12 hours (Fig. 1c). Importantly, the fact that CAS9 was not detectable in mock-control cells or upon Shield-1 withdrawal indicates that the fusion of Cas9 with the engineered FKBP12 destabilized mutant holds Cas9 expression under tight control (Fig. 1c).

Taken together, these data demonstrated that FKB12-derived destabilizing domains when fused to Cas9 enable tight, rapid and reversible control of CAS9 expression, making DD-Cas9 a suitable tool for the generation of inducible genome editing systems.

### DD-Cas9 drives robust synthetic ligand-dependent gene knockdown in mammalian cells and enables for the identification of essential genes in vitro and in vivo

To assess whether destabilized Cas9 (DD-Cas9) was competent for gene editing we targeted multiple genes controlling diverse biological process such as oncogenic transformation (EGFR), DNA damage responses (Tp53), mitochondria metabolism (CypD) and DNA replication/repair (RPA3). A549 cells were transduced with lentiviral constructs expressing DD-Cas9 and locus specific guide RNAs (Supplementary Fig. 1). Cells were treated with ligand 24 hours after lentiviral infection. At that point, cell extracts were collected after 6 days and immunoblotted to verify gene editing (Fig. 2a). The efficient formation of on-target insertion/deletion (indels) mutations in cells treated with Shield-1 was confirmed via the SURVEYOR assay (Fig. 2b). Data analysis indicated that induction of Cas9 with Shield-1 led to efficient gene editing, and resulted in a significant reduction in the expression of EGFR, CypD, RPA3 and p53 proteins.

**Figure 2.**
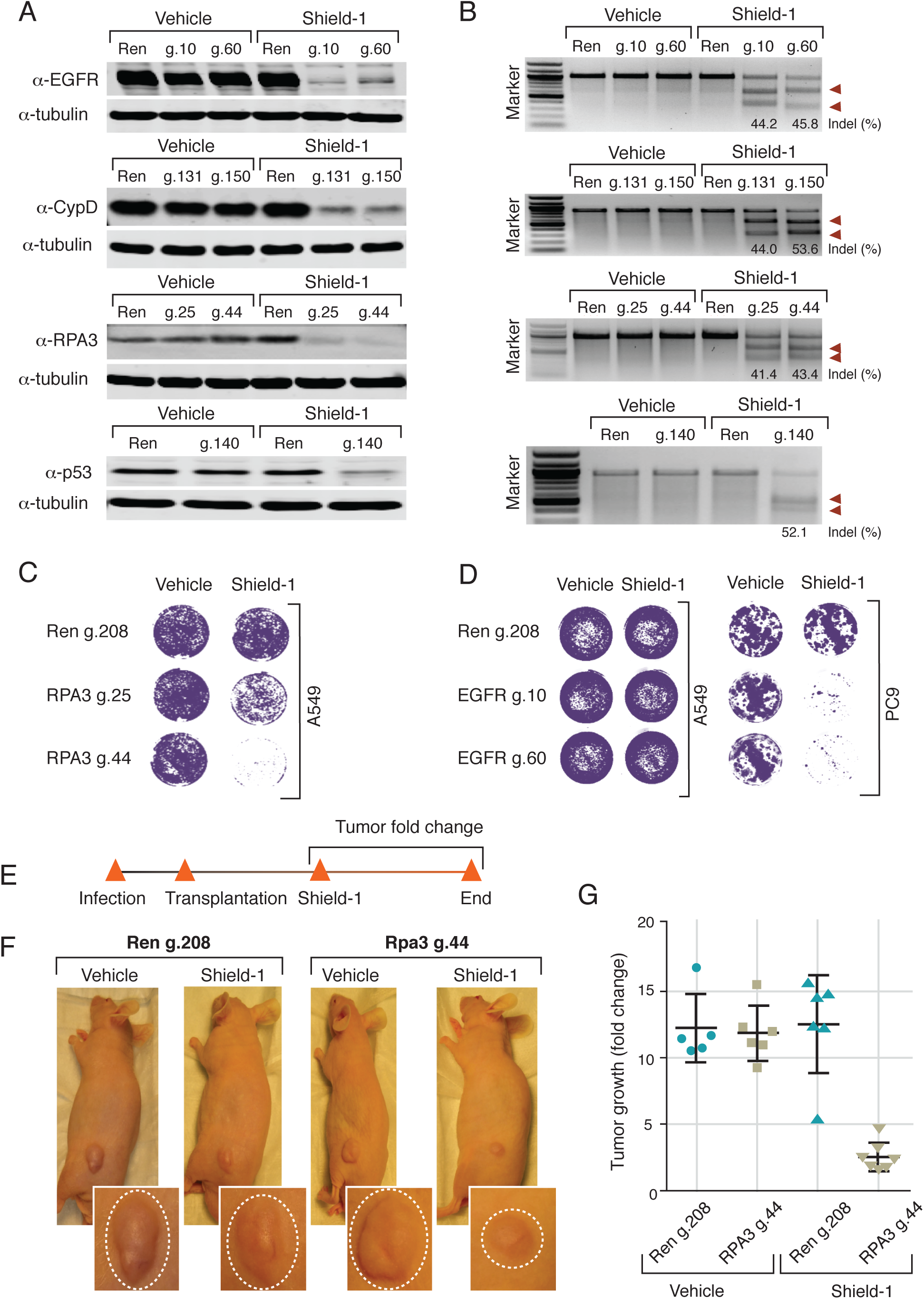
DD-Cas9 can drive robust synthetic ligands-dependent gene editing in mammalian cells and enables the identification of essential genes in vitro and in vivo. **(A)** Targeting of *EGFR, CypD, TP53 and RPA3 genes* in human A549 cells by DD-Cas9. A549 cells transduced with DD-Cas9 lentiviral vector harboring sgRNAs targeting the indicating genes. See Supplementary Fig. 1 for further information. Western blot analysis was used to assess the expression of the indicated genes before and after treatment with Sheild-1 (200 nM). Alpha-tubulin was used as a loading control. Protein expression analysis was conducted following 6 days of Shield-1 treatment. **(B)** Representative SURVEYOR assays for DD-Cas9–mediated on-target indels generation at the human *EFGR, TP53, CypD and RPA3* loci in the presence and absence of Shield-1. DNA extracts were prepared similar to what was described for the above described protein expression analysis. Arrowheads indicate expected SURVEYOR fragments. **(C)** Targeting RPA3 in A549 cells resulted in dramatic decrease in the number of viable cells following induction with Shield-1. A549 cells were transduced with the DD-Cas9 vector targeting the RPA3 locus and as control the Renilla gene. Number of cells was assessed by crystal violet staining 6 days after Shield-1 and vehicle treatment (See Supplementary Figure 3 for quantification). **(D)** Inactivation of EGFR by DD-Cas9. PC9 cells and A549 cells were transduced with DD-Cas9 vector targeting the EGFR locus. Cells were treated 24 h after infection with Shield-1 (200 nM) for 6 days. Number of cells was assessed after 6 days by crystal violet staining. (See Supplementary Figure 3 for quantification). **(E)** Workflow of the transplantable model system used in this study. **(F)** A549 cells were transduced with DD-Cas9 targeting RPA3 and as control Renilla and transplanted sub-cutaneously in immune-deficient mice. When the tumors reached an approximate size of 4-5 mm in diameter; mice were treated with Shield-1 (1 μg). Fold change in tumor growth was determined 10 days after treatment. Pictures illustrate representative tumor sizes. The chart in panel **(F)** illustrates combined quantification of tumor volume in the indicated cohorts upon Shield-1 treatment. Data represent mean ± SD and are shown relative to the negative control (Cas9 negative). p < 0.0001 (RPA3 g.44 + Shield-1 vs – Shield-1).

RPA is the heterotrimeric single-stranded DNA binding complex composed by the RPA1, 2 and 3 proteins that stabilizes replication forks by coating melted DNA (20, 21). The RPA complex also binds ssDNA at sites of DNA damage and recruits cell cycle checkpoint kinases (21, 22). Previous work showed that targeting the smallest RPA subunit, RPA3, induces degradation of all three members of the RPA complex, and associated with decreased cell proliferation and increased cell lethality (23, 24). Hence we chose to target the RPA3 locus to assess the competency of our system to study genes that are required for cell survival.

A549 cells were transduced with lentiviral constructs expressing DD-Cas9 and two independent RPA3 sgRNAs (RPA3 g.25 and g.44) and as control a Renilla sgRNA (g.208) (Supplementary Fig. 1). In the absence of Shield-1, we did not observe difference in number of cells between uninfected cells and cells infected with lentiviruses targeting RPA3 or Renilla. Yet upon treatment with Shield-1, a drastic decrease in cell number was apparent in cells infected with the lentivirus harboring the RPA3 but not the Renilla guides (Fig. 2c). Analysis of the onset of this effect indicated that just 48 hours exposure to Shield-1 compound was sufficient to induce robust decreases in cell viability (Supplementary Fig. 2 and 3).

We previously showed that the NSCLC derived PC9 cells harbor an oncogenic-EGFR mutation and are highly dependent on EGFR for their survival as opposed to the case of EGFR wild-type cells such as A549 cells (25). We took advantage of this observation to test whether DD-Cas9 could be used for the identification of genotype-driven vulnerabilities. PC9 cells were transduced with lentiviral constructs carrying the guide RNAs targeting the EGFR locus (EGFR g.10 and g.60) (Supplementary Fig. 1). As shown by the crystal violet staining and cell counting, treatment of PC9 cells with Shield-1 resulted in a prominent and rapid decrease of viable cells (Fig. 2e and Supplementary Fig. 3). This was not observed for A549 cells transduced with constructs targeting Renilla or the EGFR locus, albeit Western blot analysis of cell extracts confirmed decreased expression of EGFR in these cells.

Shield-1 compound has been shown to be not toxic and to have good pharmacodynamics and kinetic properties in mice models (14). Having shown ligand-dependent gene knockdown in *in vitro* cell culture, we reasoned that this platform could be exploited for assessing gene function *in vivo* as well. As a proof of principle, we transplanted A549 cells transduced with the DD-Cas9 vector targeting either RPA3 or Renilla into immune-compromised mice via sub-cutaneous injection (Fig. 2e). When tumors reached an approximate size of 4-5 mm in diameter, mice were treated with Shield-1 or vehicle for 4 days and tumor size was assessed for the next 10 days. We observed a drastic 5 fold decrease in tumor volume in tumors expressing RPA3 guide upon Shield-1 treatment (Fig. 2f, g).

### DD-Cas9 can be coupled to a modified fluorescent protein to generate a tractable system

An additional advantage of this system is the conditional regulation of CAS9 protein stability independent of its mRNA expression. This implies that any gene of interest can be co-expressed with destabilized Cas9 from the same promoter. Among various strategies employed to construct bi-cistronic or multi-cistronic vectors, an internal ribosomal entry site (IRES) has been widely used. More recently, a self-cleaving 2A peptide has been shown to be an alternative strategy. As a proof of principle to verify the possibility of expressing a gene of interest together with conditionally destabilized Cas9, we cloned the fluorescent protein Venus downstream of DD-Cas9 and separated them with the self-cleaving 2A peptide (P2A) (Fig. 3a). As shown in Figure 3b, expression of the modified-GFP fluorescent protein was constitutively expressed despite fusion to DD-Cas9, and was not altered by Shield-1 treatment. Similarly, the mRNA expression was independent of ligand treatment (Fig. 3b).

**Figure 3.**
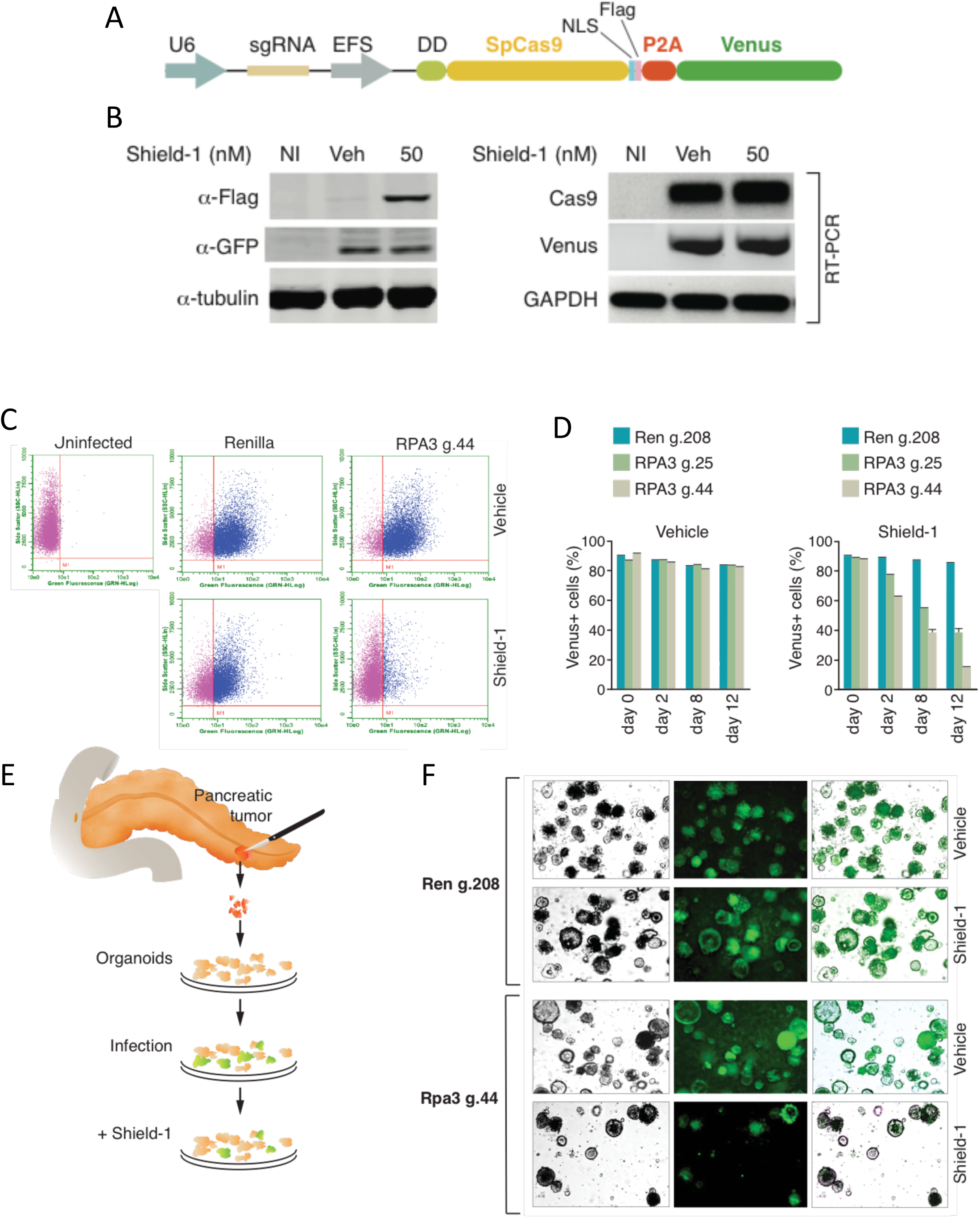
DD-Cas9 can be coupled to a modified fluorescent protein to generate a tractable system. **(A)** Schematic of the lentiviral DD-Cas9 plasmid containing U6 promoter–driven single guide RNA (sgRNA) and EFS promoter–driven DD-Cas9 pieces. Of note a FLAG tag is present at Cas9 C-terminal moiety, and 2A self-cleaving peptide (P2A) separates both DD-Cas9 and the gene of interest, in this case the modified fluorescent protein Venus. **(B)** Shield-1 independent expression of Venus. A549 cells were transduced with the DD-Cas9/P2A/Venus lentiviral vector and treated with Shield-1 for three days. Expression of Cas9 and Venus were assayed by western blot and RT-PCR. **(C)** Competition assay. A549 cells were transduced al low MOI with the DD-Cas9/P2A/Venus lentiviral vector. Cells were then treated with Shield-1 or vehicle control for 12 days and quantified by FACS. The panels represent typical FACS dot plots form A549 sorted cells. In **(D)** the chart illustrates the percentage of Venus^+^ cells during time in A549 cells transduced with the DD-Cas9/P2A/Venus lentiviral vector targeting the indicated genes upon treatment with Shield-1 and vehicle control. **(E)** Workflow to test for cell vulnerabilities in pancreatic tumor explants. **(F)** DD-Cas9/P2A/Venus could be used to infect primary human tumor derived cultures and to assess their cellular vulnerabilities. Human pancreatic cancer derived organoids were infected with the DD-Cas9/P2A/Venus lentiviral vector targeting RPA3 and as control Renilla. Organoids, cultured in 24-well plates, were treated with Shield-1 or vehicle control 72 hours post-infection. Number of Venus^+^ organoids was determined 72 hours after infection and 6 days after Shield-1 treatment. The pictures on the right panel and in Supplementary figure 4 depict representative images of the organoids.

An important consideration when developing therapeutic strategies based on decreased expression of a gene of interest by viral delivery is the potential impact on the physiology and viability of the transduced cells. This effect can be particularly relevant in longer time course experiments, where subtle differences in cell growth rates can have a major impact.

In this regard, “competition assays” have been shown to be powerful systems to be able to score for subtle cell growth changes (26). This assay is based on the competitive cell growth between transduced and non-transduced cells (27). The mixed culture obtained after gene transduction can be observed in the ratio of GFP^+^/GFP^-^ cells over time to identify minor cell growth defects (28). When compared to established methods of counting cells and viability assays (eg. MTT and ATP bioluminescence), competition assays demonstrated to provide superior sensitivity.

To verify that our platform could also be used in competition assay set-up, we infected A549 cells with the DD-Cas9/P2A/Venus lentiviral vectors targeting either RPA3 or as a negative control the Renilla gene at low MOI. This ensured the generation of Venus^+^ and Venus^-^ mixed culture (Fig. 3c). By tracking the ratio of Venus^+^ vs. Venus^-^ cells infected with the RPA3-targeting vector on a time-course of 12 days, we observed a Shield-1-dependent depletion of Venus^+^ cells (Fig. 3c, d).

The newly developed 3D “organoid” culture system allows for cultivation of both normal and patient-derived primary tumors in vitro (29). We tested whether the DD-Cas9/P2A/Venus platform could also be applicable to this culture system (Fig. 3e and Supplementary Fig. 4). Upon infection, we observed approximately 80% Venus^+^ organoids. Strikingly in organoids derived from two different tumors, after treatment with Shield-1 this fraction decreased to less than 20% for the RPA3-targeting vector (Fig. 3f and Supplementary Fig. 4). This result clearly demonstrates the utility and efficiency of the DD-Cas9 system as a screening tool in primary tumor cells.

### DD-Cas9 can be combined with tamoxifen inducible Cre (Cre-ER^T2^) for the independent conditional perturbation of genes

Recent studies indicated that CRISPR/Cas9-based approach could be used for the functional investigation of candidate genes in well-established mouse models of cancer (30, 31). In particular these studies showed that Cre-dependent somatic activation of oncogenic Kras (G12D) could be combined with CRISPR/Cas9-mediated genome editing of tumor suppressor genes by using a lentiviral-based system that delivers both the CRISPR/Cas9 system and the CRE recombinase (32).

We reasoned that by generating a bicistronic vector for the expression of destabilized Cas9 and tamoxifen inducible Cre, we could uncouple gene editing and Cre-mediated recombination. By providing temporal control of gene editing and activation/inactivation of LOX alleles, this approach will enable a more precise study of the functional interactions among genes in a variety of pre-existing mouse models derived from Cre-lox system. Especially in the context of cancer models, it could be used to dissect more precisely the role of mutations identified in onco-genomic studies during tumor onset and progression. In particular, given that cancer is a multi step process that involves the progressive accumulation of mutations, this approach could be invaluable for the identification of tumor-cell vulnerabilities during the evolution of tumors from pre-neoplastic to fully malignant diseases.

In order to achieve this, we engineered the DD-Cas9/P2A/Venus backbone to express Cre-ER^T2^ downstream of Cas9 (Fig. 4a). More specifically, we inserted an internal ribosome entry site (IRES) downstream of DD-Cas9 open reading frame (ORF) which was then coupled to tamoxifen inducible Cre (Cre-ER^T2^). In this setting, gene editing could be induced independent of Cre-mediated recombination.

**Figure 4.**
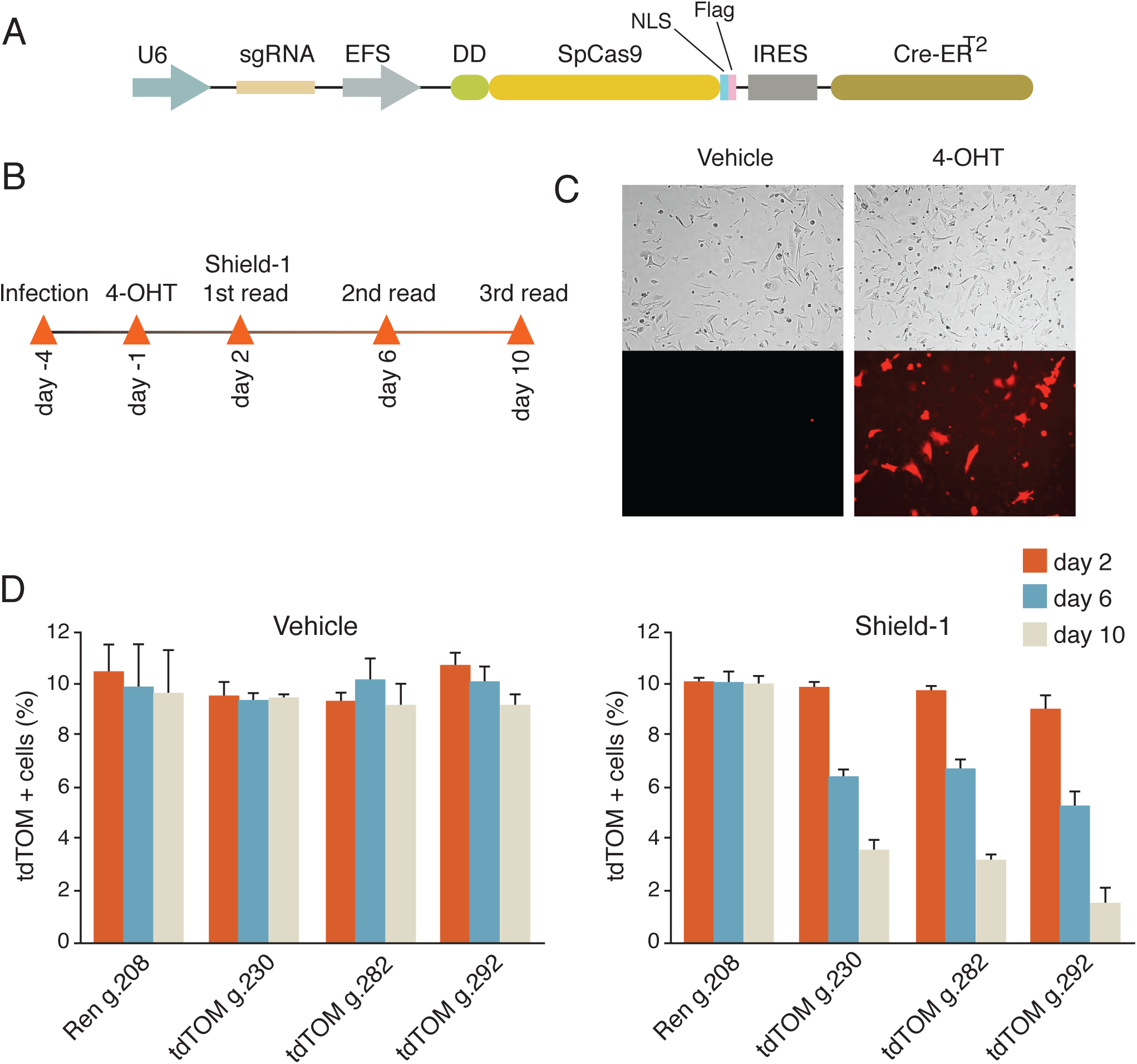
DD-Cas9 can be combined with tamoxifen inducible Cre (Cre-ER^T2^) to facilitate the study of genetic interaction in vivo in pre-existing mouse models based on Cre-lox system. **(A)** Diagram of the DD-Cas9/IRES/Cre-ER^T2^ vector utilized in this study. **(B)** Schematic for experimental design. MEF cells obtained from CAG-LSL-tdTomato mice were transduced with the indicated vectors. Induction of Cre-mediated recombination was induced upon 4-hydroxytamoxifen treatment (5 μM) for three days. To test for efficiency of gene inactivation cells were treated with Shield-1 (200 nM). **(C)** Bright-field and Fluorescence images of LSL-tdTomato cells treated with 4-hydroxytamoxifen. **(D)** The chart represents the percentage of tdTomato positive MEFs transduced with DD-Cas9/IRES/Cre-ER^T2^ lentiviruses carrying the indicated sgRNAs during time after exposure to Shield-1 and vehicle. Numbers of tdTomato positive and negative cells were determined by FACS. Each bar represents the average of three individual replicates.

To test this system, we used genetically engineered mouse lines that express the fluorescent reporter protein tdTomato in a Cre-dependent manner from the constitutive CAG promoter (tdTom^LSL^; Ai14). In this setting, following Cre-mediated deletion of the floxed stop cassette, tdTomato can be readily expressed. We derived mouse embryonic fibroblasts (MEF) and transduced them with our lentiviral vector. Three days after infection, MEFs were acutely treated with 4-hydroxytamoxifen to induce activation of Cre (Fig. 4b). We assessed expression of tdTomato by fluorescent microscopy and FACS. TdTomato positive cells were readily observed only in MEFs infected with DD-Cas9/IRES/Cre-ER^T2^ upon 4-hydroxytamoxifen treatment (Fig. 4c).

Next, to assess the gene editing efficiency of our system, we targeted the Cre-activated tdTomato knock-in reporter allele with lentiviruses expressing sgRNAs against tdTomato (sgTom) or Renilla as control. When we quantified sgRNA transduced tdTomato positive MEF cells over time, we observed a substantial reduction in tdTomato expression only in those cells treated with Shield-1 compound (Fig. 4d). Thus, this result suggests that the unique system which we have developed is a functional tool for gene editing of an endogenous allele and that Cre and Cas9 activity could be uncoupled.

In summary, these results demonstrate for the first time the potential of using destabilizing domain technology to generate a highly scalable and efficient platform for the temporal control of CRISPR/Cas9 based gene editing. Importantly, this study presents a novel approach to test for cooperativity between genetic events and identify tumor cell vulnerabilities in *in vitro* and *in vivo* models.

## DISCUSSION

The possibility of acutely inactivating genes in cells and in organism is essential to understand gene function. The most dramatic phenotype defining a gene as essential is lethality. In the context of cancer research, essential genes are of particular interest as they could provide potential therapeutic targets. Yet the fact that upon their inactivation the cells die renders them the most difficult class of genes to study.

Many different approaches have been developed and used in the past for the identification and characterization of essential genes. These methods include temperature-sensitive mutants (33); conditional deletion alleles in which a gene of interest is flanked by loxP recombination sites enabling acute gene inactivation upon expression of Cre recombinase (34); and inducible gene knockdown based on shRNA mediated silencing (35). More recently, methods to temporally control CRISPR/Cas9 based genome editing technology has also been developed. These systems are based either on DOX-induced sgRNA and Cas9 expression or on a split Cas9 system that relies on rapamycin treatment for the formation of a holo-Cas9 enzyme (11, 12).

As an alternative approach, we exploited a recently developed strategy in which a cell-permeable ligand is used in conjugation with a genetically encoded destabilizing domain (DD) to temporally control the expression of Cas9 enzyme. We showed that this system achieves a highly specific and dose-dependent conditional regulation of Cas9 expression and therefore gene editing.

In particular, we presented evidence that this system could be exploited for the identification of essential genes in standard 2D growing conditions as well as for the study of primary tumor explants such as organoids.

As the small-molecule DD ligand Shield-1 is highly permeable and nontoxic in cultured cells and animals, we also demonstrated our methods could be utilized in transplantable model system and in mice mosaic models. In particular, by coupling DD-Cas9 with tamoxifen inducible Cre, we showed that our lentiviral vector could in principle facilitate the study of genetic interaction in vivo in a variety of pre-existing mouse models of human diseases based on Cre-lox system.

Recent studies have shown that a similar lentiviral vector based on tandem expression of CAS9 and CRE recombinase could be used in vivo to rapidly evaluate human cancer genome candidates (36). Different to the existing methods, our approach enables temporal and independent control of gene editing and of Cre-mediated recombination. This, uniquely, will enable to investigate the role of putative cancer genes on tumor onset, progression and in driving tumors vulnerabilities.

In summary, our work demonstrates that the use of a conditionally destabilized Cas9 could be harnessed to generate a novel platform for the temporal control of gene editing that is very robust and easily scalable for the identification of essential genes and the interrogation of gene functional interaction in vitro and in vivo.

## METHODS

### Animals

All animal experiments were performed in accordance with National Research Council’s Guide for the Care and Use of Laboratory Animals. Protocols were approved by the Cold Spring Harbor Laboratory animal care and use committee. Male NU/NU mice at age of 6 weeks were purchased from Charles River. A549 lung cancer cells were plated and infected in vitro with lentiviruses carrying Renilla or RPA3 sgRNAs at a multiplicity of infection (MOI) of <1. Xenograft tumors of A549 cells with inducible-cas9 expression were established in the flanks of mice by subcutaneous injection of 5x10^5^ cells in 200 μL volume mixed with 1:1 dilution basement membrane matrix with biological activity (Matrigel, BD Biosciences). Five to six mice per each group were used in each experiment. Tumors were allowed to grow for 2 weeks. When tumors reached a palpable size, mice were randomly segregated into cohorts that received either four times (once every day) local peritumoral injection of Shield-1 (1 μg diluted in 100 μL PBS) or vehicle placebo on a course of four days. Tumor growth was followed for ten days using a vernier caliper (volume = ((d*short*)^*2*^ × (d*long*))/2). At the end of the experiment, mice were sacrificed. Tumors were extracted and fixed in freshly prepared 4% paraformaldehyde for 24 h.

### Cell Lines

All cell lines were obtained from American Type Culture Collection (ATCC). A549 and PC9 cells were cultured in RPMI supplemented with 5% Fetal Bovine Serum (FBS, HyClone) and 100 U/ml penicillin, and 100 μg/ml streptomycin (Gibco) at 37°C with 5% CO_2_ incubation. MEFs and Hek-293T cells were maintained in Dulbecco’s Modified Eagles Medium (DMEM) with 10% FBS and antibiotics. Virus packaging was achieved by transiently co-transfecting Hek-293T cells on 10 cm culture dish with 3 μg of the construct encoding the genes of interest and sgRNA along with 6 μg of the packaging plasmid psPAX2 and 3 μg of the envelope plasmid pMD2.G (Addgene) using 30 μL of the Lipofectamine2000 reagent (Life Technologies). Ten mL viral particles were collected after 48 h of transfection by clarifying the supernatant through 0.45 μm filter membrane (GE Healthcare). Virus transduction was optimized in order to achieve low MOI transduction. Typically, 500 μL fresh virus particles from 10 mL stock were used to infect 1x10^6^ cells on a 10 cm culture dish in 10 mL total volume of culture medium. Virus aliquots were stored at -80°C. Shield-1, obtained from Cheminpharma, was solubilized in pure ethanol and was added to culture media with given concentrations.

### Organoids

Methods to establish and propagate human organoids cultures were previously described. Organoid infections were performed as described. In brief, organoids were grown into a 24-well culture plate for two days before infection. One well per infection was used. The day of infection, organoids were dissociated into small fragments by first triturating them in media through a fire-polished glass pipette, and then by a 5 min enzymatic digestion with TrypLE (Life Technologies) at 37°C. The resulting small cell clusters were then resuspended with low MOI lentivirus and spinoculated at 600 RCF for 1 hr at room temperature.

### Plasmids

Vectors used in this study were engineered based on the LentiCRISPR V2 (Addgene) backbone. P2A-linked puromycin located downstream of the SpCas9 was substituted either Venus, a modified version of Green Fluorescent Protein (GFP) for easy visualization of viral transduction efficiency as well as competition assay. For Cre-ER^T2 co-expression^

High-affinity ligand dependent destabilization domain (DD) of a mutant FKBP12 protein (F36V, L106P) was inserted at the N-terminus of SpCas9. BsmBI digestion site in the DD-sequence was replaced with silent mutations by site-directed mutagenesis.

### Western Blot

Protein samples were isolated by resuspending cell pellets in RIPA buffer (50 mM Tris-HCl at pH 7.6, 150 mM NaCl, 1% NP-40, 0.5% Na deoxycholate, 0.1% SDS). After removal of the debris, samples were quantified with colorimetric BCA kit (Pierce). 20 μg total proteins were electrophoresed on 6-12% gradient gels and wet-transferred to nitrocellulose membranes. After 1 h blocking with 5% nonfat dry milk in 1X TBS, 0.1% Tween20 at room temperature, membranes were incubated with antibodies diluted in 2% w/v BSA as follows; Flag [M2] mouse mAb (1:1000, Sigma) to detect DD-SpCas9 expression, α-tubulin [DM1A] mouse mAb (1:20000, Millipore) as equal loading control, GFP [C163] mouse mAb (1:5000, Invitrogen) to detect Venus expression, p53 [DO-1] mouse mAb (1:1000, Millipore), CypD [E11AE12BD4] mouse mAb (1:5000, Abcam), RPA14 [11.1] mouse mAb (1:1000, Abcam) and DD monoclonal Ab (1:1000, Clontech). All incubations were performed overnight at 4°C. Membranes were rinsed thoroughly with 1X TBS-T and then incubated with species-specific fluorescently labeled secondary antibodies (1:5000, LICOR). Western blots were eventually imaged on near-IR fluorescence scanner (Odyssey Imaging System, LICOR).

### RNA isolation and RT-PCR

Cells were rinsed twice and harvested with ice cold PBS. Pellets were lysed in 1 ml Trizol (Invitrogen) and RNA was extracted according to the manufacturer’s instructions. Contaminating DNA was removed by RNase-free DNase (Promega) treatment for 30 min at 37°C. cDNA was prepared from 2 μg total RNA using ImProm-II Reverse Transcription System (Promega) with 16mer oligo(dT). Semi-quantitative RT-PCR detection was performed using *Taq* DNA Polymerase with standard *Taq* buffer (NEB) using primers specific to SpCas9, GFP and GAPDH.

### SURVEYOR assay

Cells were infected with lentivirus and incubated at 37C for indicated times as described. Genomic DNA was extracted using mini-spin columns following the manufacturer’s instructions (Macherey Nagel). In brief, cell pellets were resuspended in Lysis Buffer and incubated at 56C for 3 h and 70C for 10 min. SpCas9-induced mutations were detected using the SURVEYOR Mutation Detection Kit (Transgenomic/IDT). Approximately 900 bp region surrounding the CRISPR mutation target site was PCR-amplified using Phusion High-Fidelity DNA Polymerase (NEB). Single band PCR products were purified using USB PrepEase Gel Extraction kit (Affymetrix). 450 ng purified PCR fragments mixed with 2 μL 10X Taq DNA Polymerase PCR buffer in a total volume of 20 μL were subjected to a series of melt and reanneal temperature cycles with gradual increments lowered in each thermal cycle. SURVEYOR nuclease and SURVEYOR enhancer were then added to the reaction mixture in order to selectively digest heteroduplex DNA substrates. Digested products were run and visualized on a 1.2% agarose gel. Band intensities were quantified using ImageJ software. Indel percentage was calculated using the formula 100 × (1 – (1 – (*b* + *c*)/(*a + b + c*))^1/2^), where *a* stands for the integrated intensity of the undigested product and *b* and *c* are the integrated intensities of each cleaved PCR product.

### Crystal Violet Staining

Cells infected with lentiviruses were plated in equal number in 12 well plates (BD falcon) with expression of Cas9 stabilized using 200 nM of Shield-1 and incubated at 37°C for indicated times as described. Cells were rinsed twice with PBS to eliminate the floating cells and fixed in 4% Formaldehyde in PBS (V/V) for 10-15 minutes. Fixed cells were stained by staining solution (0.1% Crystal violet in 10% ethanol) for 20 minutes. Staining solution was aspirated from the wells and cells were washed with water thrice in order to remove extra stain. Stained cells were air dried and imaged using Odyssey Imaging System (LICOR). To quantify the relative cell numbers, cells were destained with 10% Acetic acid and absorbance of destained solution was measured at 590 nm at appropriate dilutions.

---

**Supplementary Figure 1:**
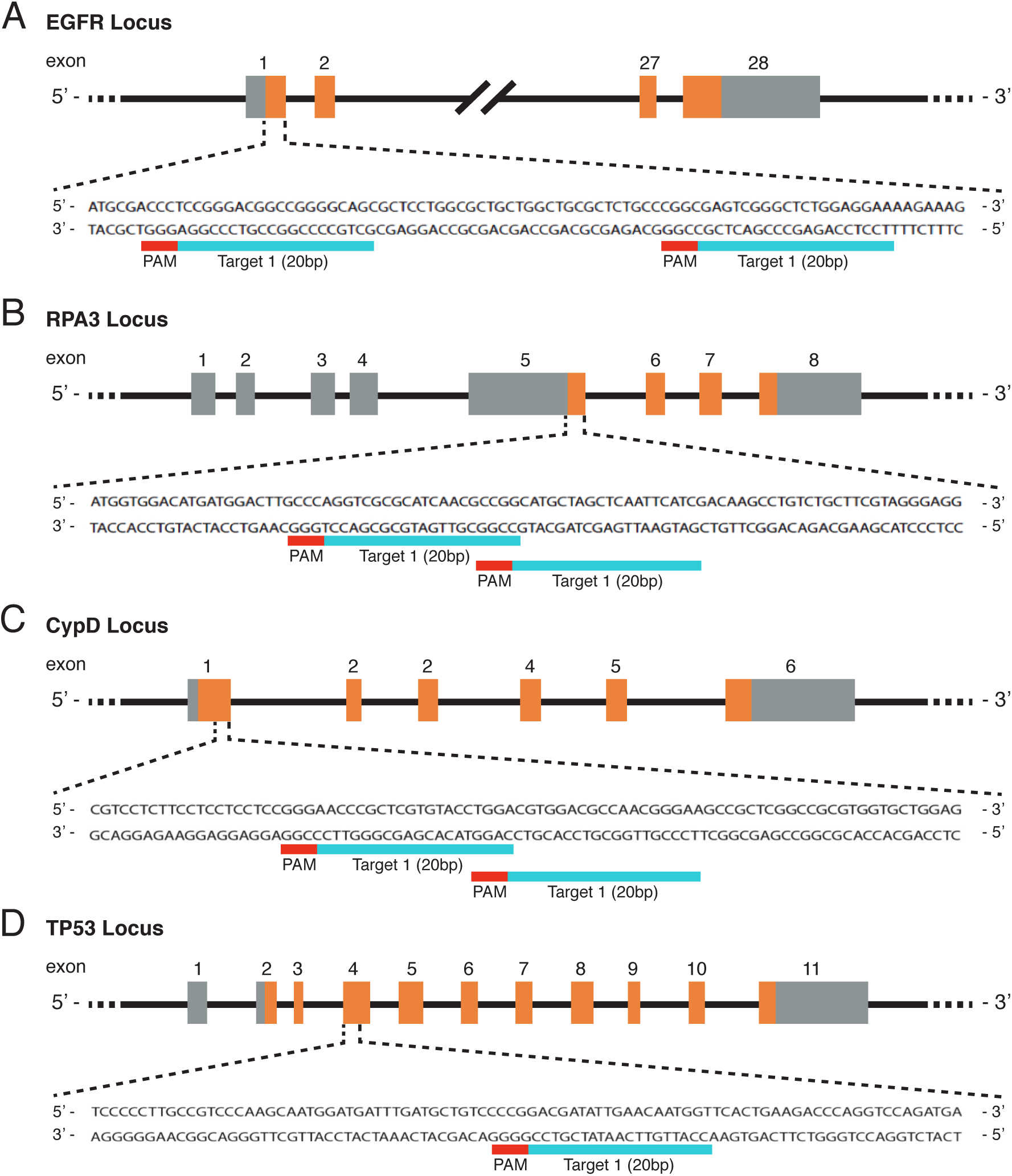
Schematic of location of small guide RNAs on EGFR, RPA3, CypD and TP53 genes loci. Guide RNA predictions were performed using Optimized CRISPR design tool from MIT.

**Supplementary Figure 2.**
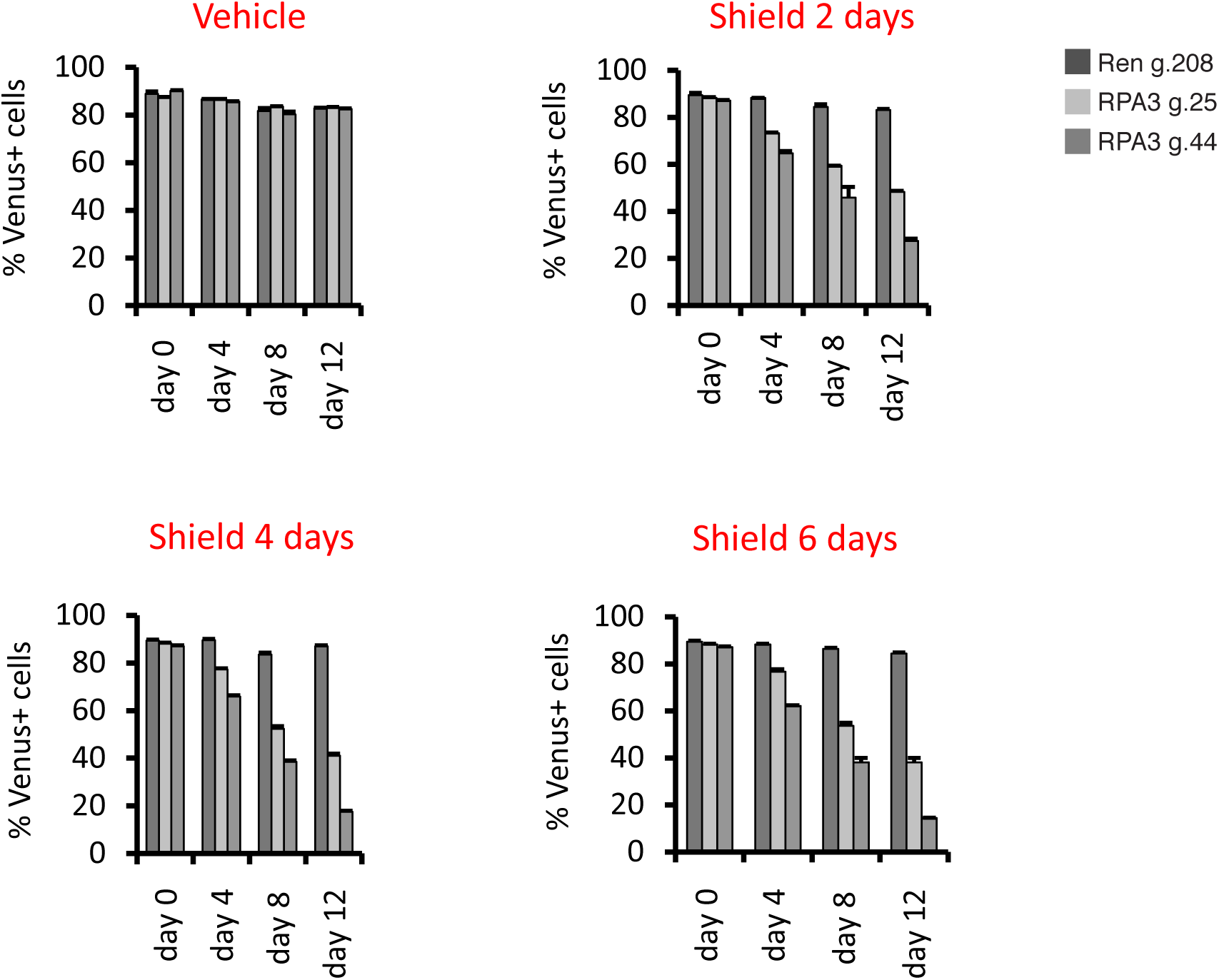
DD-Cas9 can be coupled to a modified fluorescent protein to generate a tractable system. A549 cells were transduced with the DD-Cas9/P2A/Venus lentiviral vector. Cells were then treated with Shield-1 or vehicle control and quantified by FACS. The panels illustrate the percentage of Venus+ cells during time in A549 cells transduced with the DD-Cas9/P2A/Venus lentiviral vector targeting the RPA3 gene with two individual sgRNAs. Upon treatment with Shield-1 and vehicle control for different time points, it is evident that only 2 days of Shield-1 treatment was sufficient to induce similar changes with 4 and 6 days continuous ligand treatment.

**Supplementary Figure 3:**
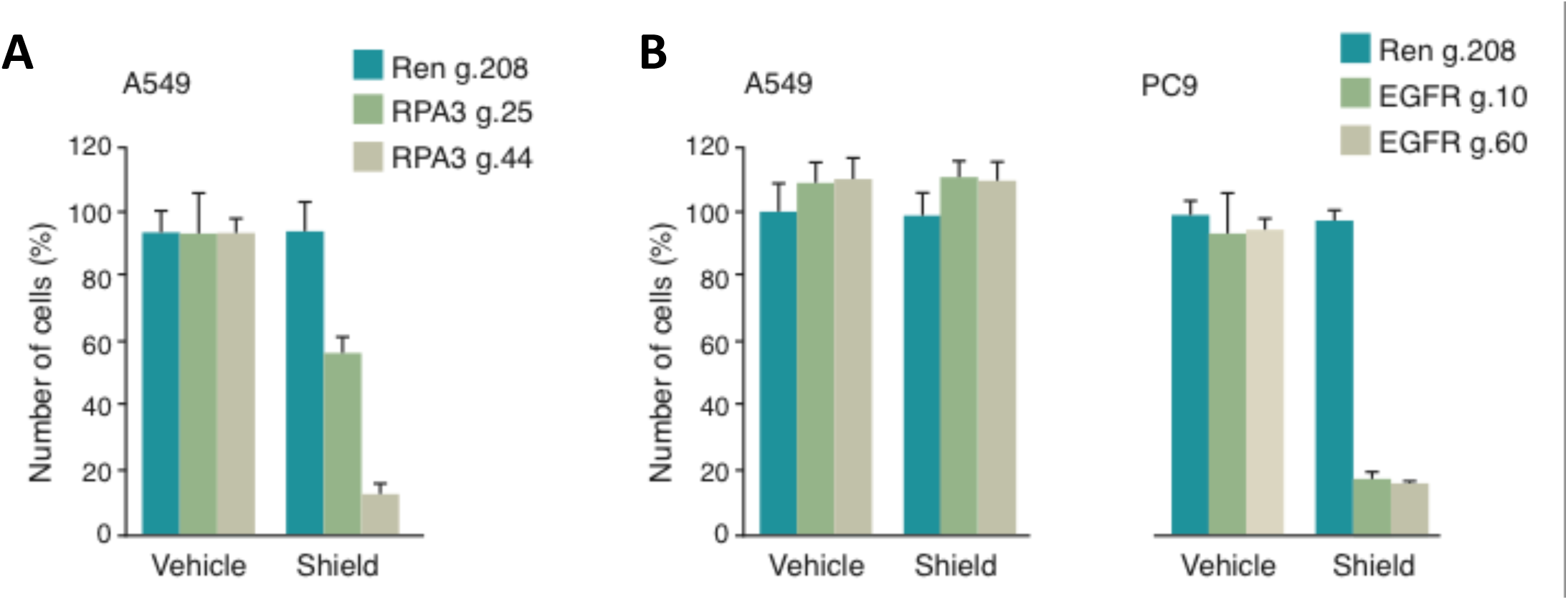
**(A)** Targeting RPA3 in A549 cells resulted in dramatic decrease in the number of viable cells following induction with Shield-1. A549 cells were transduced with the DD-Cas9 vector targeting the RPA3 locus and as control the Renilla gene. Number of cells was assessed by crystal violet staining 6 days aGer Shield-1 and vehicle treatment (Figure 2C). The data in the chart represent mean ± SD of the number of A549 cells upon treatment with Shield-1 and vehicle control and are shown relative to uninfected cells (n=4). **(B)** Inactivation of EGFR by DD-Cas9. PC9 cells and A549 cells were transduced with DD-Cas9 vector targeting the EGFR locus. Cells were treated 24 h aGer infection with Shield-1 (200 nM) for 6 days. Number of cells was assessed aGer 6 days by crystal violet staining (Figure 2D). The data in the charts represent quantification of Panel E as means ± SD upon treatment with Shield-1 and vehicle control and are shown relative to uninfected cells (n=4).

**Supplementary Figure 4.**
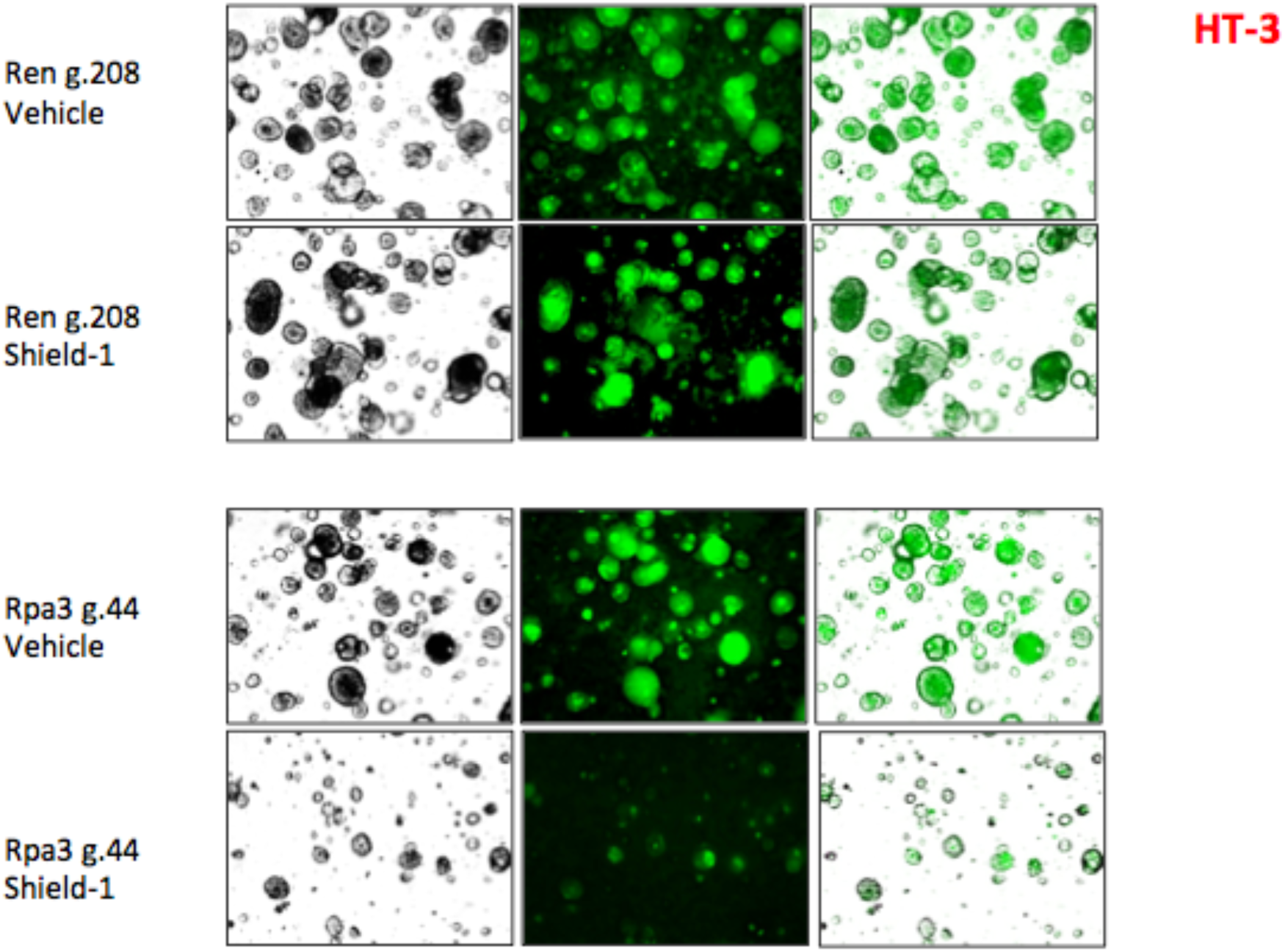
DD-Cas9 can be coupled to a modified fluorescent protein to generate a tractable system. A549 cells were transduced al low MOI with DD-Cas9/P2A/Venus lentiviral vector. Cells were then treated with Shield-1 or vehicle control for 12 days and quantified by FACS. The panels illustrate with percentage of Venus^+^ cells during time in A549 cells transduced with the DD-Cas9/P2A/Venus lentiviral vector targeting the RPA3 gane with two individual sgRNAs. Upon treatment with Shield-1 and vehicle control for different time points, it is evident that only 2 days of Shield-1 treatment was sufficient to induce similar changes with 4 and 6 days continuous ligand treatment.

